# FCRL5 is a fucose-sensitive IgG-Fc receptor with binding properties distinct from classical Fcγ receptors

**DOI:** 10.64898/2026.07.01.735886

**Authors:** Nick van der Hoeven, Miles D Holborough-Kerkvliet, Yixiang Bao, Arthur EH Bentlage, Pleuni de Heer-Ooijevaar, Ninotska IL Derksen, Timon Damelang, Bart-Jan de Kreuk, Aran F. Labrijn, Gestur Vidarsson, Theo Rispens

## Abstract

Fc receptor-like protein 5 (FCRL5) is a low-affinity IgG receptor expressed on B cells, with emerging therapeutic relevance due to its expression on multiple myeloma cells, and a potential role in regulating B cell responses. Previous reports on the FCRL5-IgG interaction vary widely in reported affinities, binding differences across IgG subclasses, and molecular requirements for maximal binding. Furthermore, the impact of Fc-engineering strategies, as used in (therapeutic) monoclonal antibodies, remains poorly understood. Here, we provide a comprehensive biochemical analysis of the FCRL5-IgG interaction. We demonstrate that FCRL5 is a true IgG Fc-receptor, binding with very low affinity (60-80 µM). FCRL5 binds IgG in a manner involving primarily the two N-terminal domains of FCRL5, and the third domain for maximal binding, but with distinct essential residues in the IgG Fc-tail. Surface plasmon resonance analysis of the binding of FCRL5 to the various IgG subclasses revealed a preference for IgG1 and IgG4. Interestingly, various Fc-engineered IgG variants commonly used for silencing or enhancing of Fc receptor binding do not impact FCRL5 binding. Screening the binding of a set of IgG antibodies carrying defined sets of Fc-mutations to FCRL5 revealed E293 as a key binding determinant and led to the discovery of E293R as a mutation that selectively abrogates FCRL5 binding while preserving binding to other classical FcγRs. Lastly, we show that FCRL5 has considerable preference for binding afucosylated IgG. Together, our results define the essential characteristics of the IgG-FCRL5 interaction and demonstrate the potential of both naturally occurring IgG variants as well as therapeutically explored bioengineered IgG formats to differentially engage FCRL5.

## Introduction

B cells play a central role in humoral immunity through the production of antibodies (Abs), specialized effector molecules that are not only able to neutralize pathogens, but also bridge adaptive and innate immunity via the antigen-specific recruitment of innate immune cells^1^.

In serum, the predominant antibody isotype is immunoglobulin G (IgG)^2,3^. In part, IgG effector functions are mediated through interactions between the IgG Fc-region and IgG Fc-receptors (FcγRs). Activatory receptors such as FcγRI, FcγRIIA and FcγRIIIA are expressed on innate immune cells such as macrophages, neutrophils and NK cells, while the sole inhibitory FcγR, FcγRIIB, is expressed on B cells.

A less-well known family of receptors that shares homology with the FcγR family, is the Fc receptor-like (FCRL) family^4^. FCRL5, the largest FCRL, is expressed on B cells and has gained recent attention for being one of the defining marker of double-negative 2 (DN2) B cells, a subset of B cells that are less associated with canonical germinal center B cell responses and more associated with early and extrafollicular B cell responses^5,6^. DN2 B cells were initially found to be increased in chronic infections and various autoimmune diseases, although more recent reports indicate that DN2 B cells are a physiologic part of protective B cell responses^6,7^. Moreover, FCRL5 is overexpressed in B cell-related hematological malignancies such as multiple myeloma, and may play a key role in myelomagenesis^8,9^. Therefore, FCRL5-directed CAR-T cells^10,11^, FCRL5-directed monoclonal Ab-drug-conjugates (ADCs)^8^ and FCRL5/CD3-directed bispecific Abs^12^ are currently being evaluated in preclinical and clinical studies.

FCRL5 consists of nine Ig-like domains and carries two intracellular inhibitory motifs and one activating motif (**Figure 1A**), suggesting that FCRL5 may affect B cell biology in a context-dependent manner^4^. To this point, it has been reported that FCRL5 functions as an inhibitory receptor that requires B cell receptor (BCR) co-engagement^13^, while a single report revealed CD21 co-engagement as a switch to an activatory role of FCRL5^14^. FCRL5 has been shown to bind to IgG with relatively low affinity with its first 3 domains^15–17^. These data suggest a potential role for FCRL5 in regulating antigen-specific B cell responses via the engagement of IgG immune complexes. Feasibly, FCRL5, without CD21 co-expression may play a similar role as the inhibitory FcγRIIB^18^, whereas FCRL5/CD21 co-expression may allow for B cell activation.

**Figure 1:**
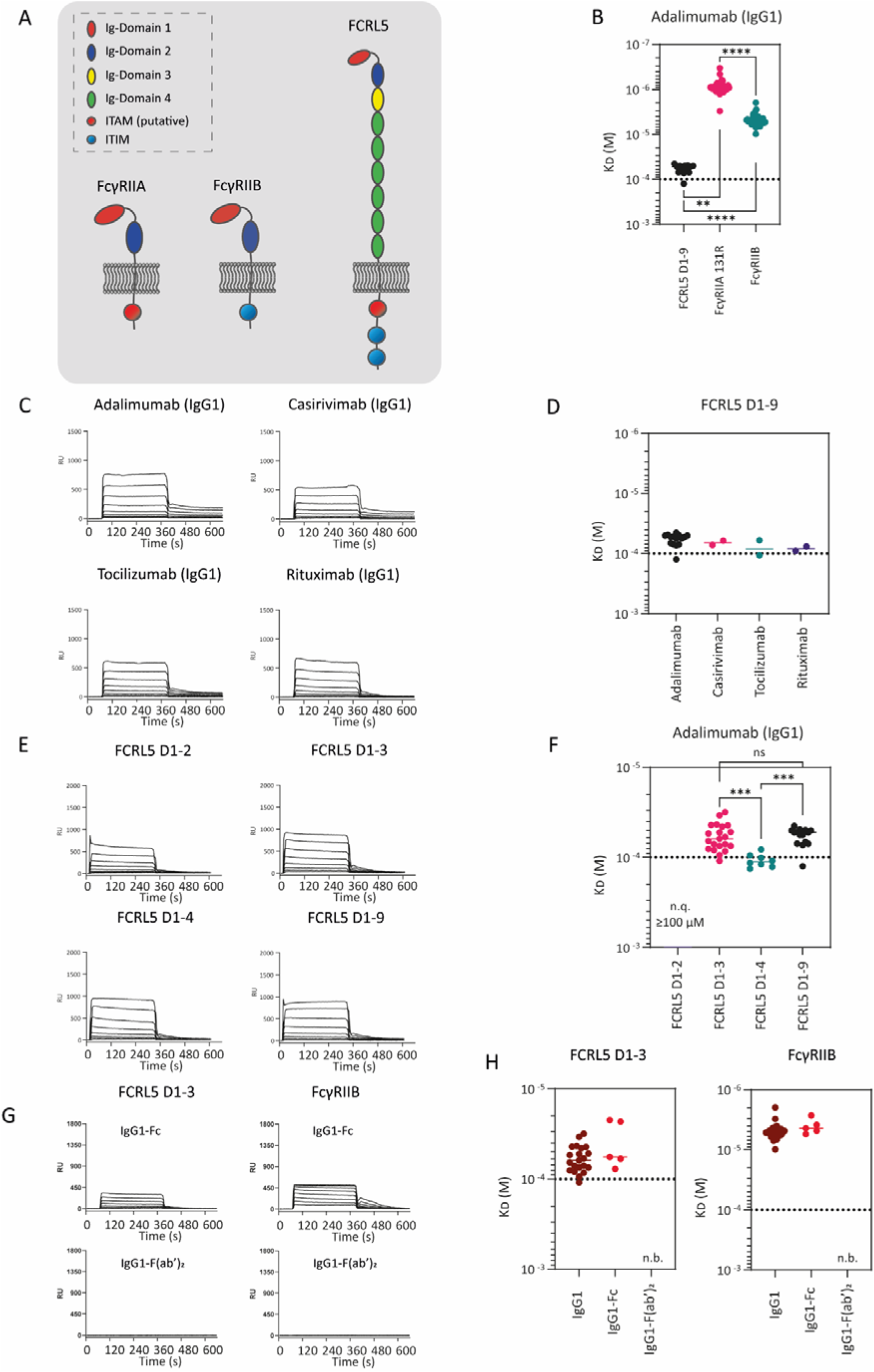
Biochemical analysis of the FCRL5-IgG interaction by surface plasmon resonance (SPR) (A) Visual representation of FcγRIIA, FcγRIIB and FCRL5. (B) Graph depicting the calculated affinities of FCRL5 D1-9, FcγRIIA 131R, FcγRIIB for adalimumab (IgG1). ** indicates a p-value <0.01, **** indicates a p-value <0.0001, determined using Kruskal-Wallis test corrected for multiple comparisons with Dunn’s correction. (C) Representative SPR sensorgrams of IgG1 mAbs (adalimumab, tocilizumab, rituximab and casirivimab) binding to FCRL5 D1-D9. (D) Graph depicting the calculated affinities of IgG1 mAbs (adalimumab, tocilizumab, rituximab and casirivimab) for FCRL5 D1-D9. (E) Representative SPR sensorgrams of FCRL5 D1-2, D1-3, D1-4 and D1-9 to adalimumab (IgG1). (F) Graph depicting the calculated affinities of FCRL5 D1-2, D1-3, D1-4 and D1-9 for adalimumab (IgG1). *** indicates a p-value <0.001, determined using Kruskal-Wallis test corrected for multiple comparisons with Dunn’s correction. (G) Representative SPR sensorgrams of IgG1 (adalimumab), IgG1-Fc and IgG1-F(ab’)_2_ (adalimumab) binding to FCRL5 D1-3 and FcγRIIB. (H) Graph depicting the calculated affinities for IgG1 (adalimumab), IgG1-Fc and IgG1-F(ab’)_2_ (adalimumab) for FCRL5 D1-3 and FcγRIIB. The dotted line represents the highest concentration of analyte measured (100 µM). Calculated values below this line are less reliable. Not quantifiable (n.q.) indicates binding that was too low for affinity calculations. No binding (n.b.) indicates no binding was observed.

It was previously reported that unlike classical FcγRs, FCRL5 engages IgG in a manner that requires both the IgG-Fc and IgG-F(ab’)_2_ for maximal binding^16^. A recent study showed a several-fold increase in affinity of FCRL5 for intact IgG compared to IgG-Fc using surface plasmon resonance (SPR), potentially supporting this notion^17^. Conversely, the Cryo-EM structure of FCRL5 and an IgM tailpiece-hexamerized IgG molecule that was featured in the study did not show any interactions between the IgG-F(ab’) and FCRL5^17^. Conflicting data exist in literature regarding the finer details of the FCRL5-IgG interaction, such as the strength of interaction to different IgG subclasses, and the reported contribution of F(ab’) to the interaction. Furthermore, data on the effects of Fc- and glycoengineered IgG variants are missing, which are relevant for the development of therapeutic mAbs. Namely, many therapeutic mAbs rely on IgG Fc-binding receptors for their mode of action, such as rituximab, obinutuzumab and ofatumumab, or in the case of blocking mAbs, require avoidance of IgG Fc-binding receptors^19^. To this end, many IgG Fc-engineered Ab formats have been developed to specifically enhance, or silence binding to specific FcγRs or the neonatal Fc receptor (FcRn)^20^. Furthermore, due to the immunologically relevant effects of increased Fc-glycan galactosylation and decreased fucosylation, which increase complement-activation and increase FcγRIII-binding^21–23^, respectively, the effects of glycoengineering on FCRL5 binding are another important aspect to investigate.

Here, we provide a detailed biochemical analysis of FCRL5 and its interactions with IgG. Using a series of well-characterized materials, we performed a thorough analysis of the binding characteristics of variants of FCRL5 as well as IgG, including different human subclasses, glycovariants, as well as a series of common, bioengineered FcγR- and FcRn-silencing or enhancing Fc formats. In addition, we exploited a panel of Fc mutants previously used to map rheumatoid factor binding^24^.

## Results

### FCRL5 is a very low-affinity IgG Fc receptor

To characterize the FCRL5-IgG interaction, we used recombinant and site-specifically C-terminally biotinylated FCRL5 from both commercial and in-house sources. HP-SEC revealed considerable aggregation of FCRL5 in both variants (**Supplementary Figure 1A).** We therefore fractionated the commercial FCRL5 and collected the monomeric fraction (from here on termed “FCRL5-D1-9”) by HP-SEC for subsequent use. Next, we employed SPR to assess the affinity of FCRL5 for IgG1. To this end, FCRL5-D1-9 was captured on a StrepA sensor chip in a titrated-manner and assessed for binding to the different therapeutic IgG1 mAb, starting at 100 µM (**Figure 1B**). FCRL5 D1-9 bound to adalimumab with affinities between 60-80 µM, affinities 1 to 2 orders of magnitude lower than FcγRIIB and FcγRIIA, respectively, which were taken along as both positive controls and comparators for IgG binding (**Figure 1C**). Furthermore, casirivimab, tocilizumab and rituximab, various other IgG1 therapeutical mAbs showed similar affinities to FCRL5 (**Figure 1D**).

We aimed to pinpoint the FCRL5 domains relevant for IgG binding. Therefore, we recombinantly produced truncated FCRL5 variants consisting of the first two N-terminal domains (FCRL5 D1-2), the first three N-terminal domains (FCRL5 D1-3) and the first four N-terminal domains (FCRL5 D1-4). Interestingly, HP-SEC analysis showed no or only minimal aggregation of the truncated FCRL5 variants (**Supplementary Figure 1B**). SPR analysis of IgG binding to the FCRL5 variants indicated that considerable binding of IgG1 was observed with just the FCRL5 D1-2 fragment. Inclusion of domain three fully recapitulated binding of FCRL5 D1-9 (**Figure 1E**). No further enhancement of binding strength was observed by adding D4. Hence, due to the similar binding properties of FCRL5 D1-3 and FCRL5 D1-9, and the absence of aggregates, FCRL5 D1-3 was used for subsequent experiments.

Previous reports have suggested that FCRL5 requires intact IgG (i.e. both Fc and F(ab’)_2_) for maximal binding^16^. To probe the relative importance of the IgG Fc and the IgG F(ab’)_2_ in FCRL5 interactions, we assessed the binding of intact IgG1 (adalimumab) to recombinantly-produced IgG1 Fc and proteolytically-generated IgG1-F(ab’)_2_ (adalimumab) to FCRL5 D1-3 by SPR (**Figure 1F**). Again, FcγRIIB was taken along as a positive control and comparator for IgG binding. Our data show that, similarly to FcγRIIB, FCRL5 binds intact IgG1 with the same affinity as IgG1 Fc of ∼60 uM. Moreover, no binding of IgG1 F(ab’)_2_ was observed to FCRL5, as for FcγRIIB. Together, these data indicate that FCRL5 is a bona fide IgG Fc-receptor, similarly to classical FcγRs, that requires D1-3 for maximal binding.

### FCRL5 preferentially binds IgG1 and IgG4

Due to the low affinity of FCRL5 for IgG, large amounts of antibody are required to accurately calculate the affinities in SPR experiments (i.e. ≥ KD). Therefore, we set up an ELISA assay that assesses effective multivalent IgG binding to FCRL5, thereby requiring substantially less material (**Figure 2A**). To this end, this ELISA employs the dimerization of IgG using a proteolytically-generated murine human kappa light chain-specific F(ab’)_2_ that was shown to specifically form IgG-dimers and not higher-order complexes^25^. We assessed the binding of all IgG subclasses (IgG1-4) to FCRL5 D1-3 (**Figure 2B**). Various IgG1 mAbs, i.e., clone 7D8, adalimumab and rituximab showed comparable binding profiles to each other as well as to IgG4 mAbs nivolumab and natalizumab, whereas sutimlimab showed an almost three-fold reduced binding profile compared to the other IgG4 mAbs. The IgG2 mAb panitumumab and an in-house produced IgG2 version of golimumab were modestly reduced in binding to FCRL5. IgG3 mAbs of two different allotypes with hinge region exon 1 or hinge region exons 1-3, respectively, IgG3*01-7D8 and IgG3*03-7D8, showed a 5- to 10-fold reduction in binding to FCRL5 D1-3. Binding of these two IgG3 allotypes to FcγRIIB was not reduced and comparable to IgG1 and IgG4 (**Supplementary Figure 2A**)

**Figure 2:**
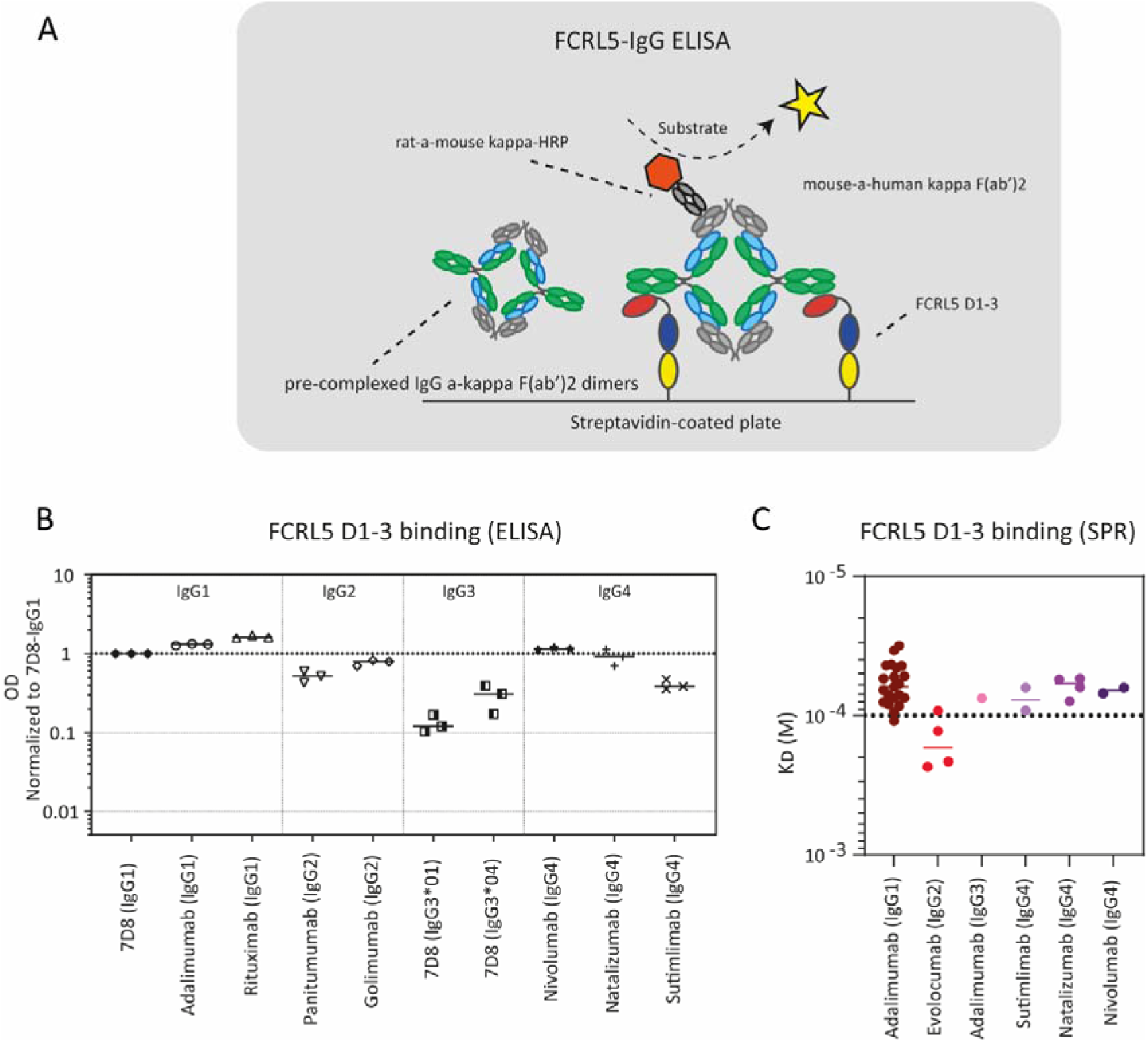
Avidity-driven ELISA setup allows for screening and relative quantification of binding of IgG subclasses to FCRL5 D1-D3. (A) Visual representation of the ELISA setup. In short, biotinylated FCRL5 D1-3 is captured on a streptavidin-coated chip. The divalent avidity provided by pre-complexed IgG-a-kappa light chain F(ab’)_2_ dimers allows for FCRL5 D1-3 binding at manageable concentrations. HRP-conjugated a-mouse kappa light chain allows for complex detection. (B) Binding of various IgG1-4 mAbs to FCRL5 D1-3 (n=3). ODs are normalized to 7D8-IgG1. (C) Graph depicting the calculated affinities of FCRL5 D1-3 for various IgG1-4 mAbs. The dotted line represents the highest concentration of analyte measured (100 µM).

To substantiate our findings, we assessed the binding of various IgG1-4 mAbs to FCRL5 D1-3 by SPR (**Figure 2C**). Evolocumab, an IgG2 mAb, showed a reduced affinity (∼167 µM) for FCRL5 D1-3, in line with ELISA results. On the other hand, despite the lower binding in ELISA, IgG3 exhibited affinities for FCRL5 comparable to IgG1 and IgG4. However, HP-SEC analysis indicated IgG3 to contain a fraction of aggregates (which we were unable to fully remove), consistent with IgG3’s reported tendency for self-association^26^, and possibly leading to an overestimated affinity. IgG4 mAbs exhibited affinities similar to those observed for IgG1. Taken together, these data show that FCRL5 preferentially binds IgG1 and IgG4 and slightly disfavors IgG2 binding, whereas for IgG3 our data tentatively suggests lower to comparable binding relative to IgG1.

### FCRL5-IgG binding is differentially affected by Fc-silenced and Fc-enhanced variants

Given the importance of Fc receptor interactions in antibody biology, we assessed the impact of established Fc-silencing and Fc-enhancing IgG mutations on FCRL5 binding. To this end, we screened a set of FcγR- and FcRn-silenced and -enhanced IgG variants for their binding to FCRL5 by ELISA. FcγRIIB was taken along as comparator. As expected, all FcγR-silenced variants abrogated binding to FcγRIIB while the FcRn-silenced variant had no effect (**Figure 3A**). In contrast, most FcγR-silenced variations barely affected binding to FCRL5. Of note, LALAPG (L234A, L235A and P329G) mutations had no significant influence on FCRL5 binding. Similarly, deletion of G236 (ΔG236) alone or in combination (i.e. in combination with E233P-L234V-L235A) (from here on referred to as “IgG1lh2”) did not affect FCRL5 binding. Only the N297A (i.e. Fc-glycan knockout) showed a strong reduction in binding to FCRL5. Consistent with these results, SPR analysis revealed that IgG1-LALAPG maintained its affinity for FCRL5, while addition of the N297A mutation (i.e. double-mutant IgG1-LALAPG/N297A) strongly impaired binding to FCRL5 **(Figure 3B, C).** In contrast, both IgG1 PGLALA and IgG1-LALAPG/N297A completely abrogated binding to FcγRIIA 131R and FcγRIIB, as previously reported^20^.

**Figure 3:**
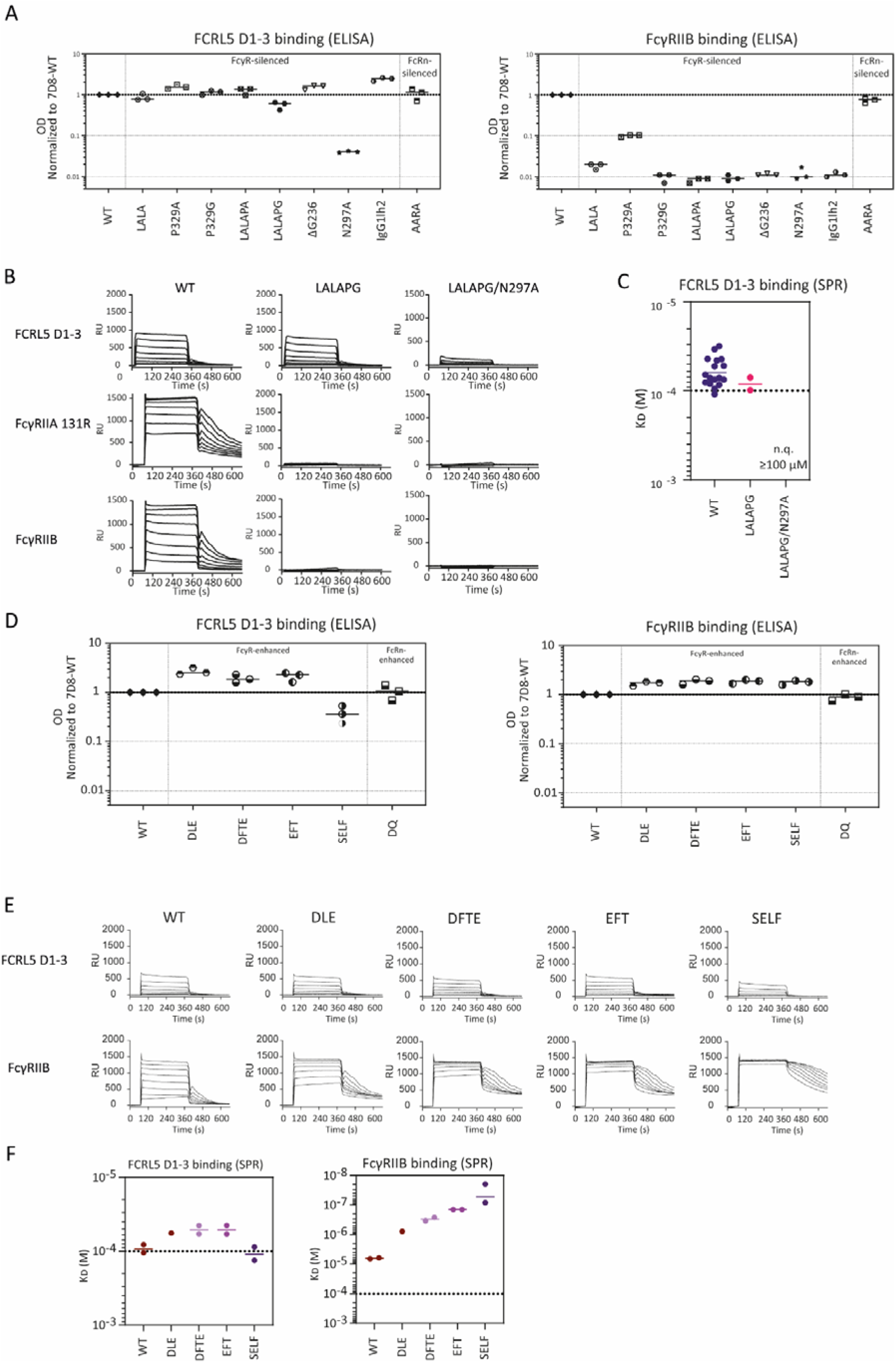
Binding of Fc-engineered IgG1 variants to FCRL5 D1-D3. (A) Binding of various Fc-silenced mAbs to FCRL5 D1-3 and FcγRIIB (n=3). ODs are normalized to 7D8-IgG1. (B) Representative SPR sensorgrams of adalimumab IgG1-WT, IgG1-LALAPG and IgG1-LALAPG/N297A binding to FCRL5 D1-3, FcγRIIA 131R and FcγRIIB. (C) Graph depicting the calculated affinities of FCRL5 D1-3 for adalimumab IgG1-WT, IgG1-LALAPG and IgG1-LALAPG/N297A. (D) Binding of various Fc-enhanced mAbs to FCRL5 D1-3 and FcγRIIB (n=3). ODs are normalized to 7D8-IgG1. (E) Representative SPR sensorgrams of various Fc-enhanced mAbs to FCRL5 D1-3 and FcγRIIB. (F) Graph depicting the calculated affinities of FCRL5 D1-3 for various Fc-enhanced mAbs. The dotted line represents the highest concentration of analyte measured (100 µM). Not quantifiable (n.q.) indicates binding that was too low for affinity calculations.

Fc-enhanced variants revealed different effects on FCRL5 binding (**Figure 3D**). Namely, the FcRn-enhanced variant DQ (T256D-T307Q) had no effect on either FcγRIIB or FCRL5 binding while, the FcγR-enhanced variants DLE (S239D, A330L and I332E), DFTE (S239D, S267E, H268F and S324T) and EFT (S267E, H268F and S324T) showed increased binding to both FcγRIIB and FCRL5. In contrast, the FcγRIIB-enhancing variant S267E-L328F displayed reduced binding to FCRL5. These findings were confirmed by SPR, although the observed changes in affinity for FCRL5 were modest and increased 1.8-fold at most, for IgG1-DFTE and IgG1-EFT **(Figure 3E, F).**

These data taken together, highlight a distinct binding mechanism of FCRL5 compared to those of FcγRIIB, as besides IgG1-N297A, most common Fc-silencing IgG variants did not abrogate FCRL5-binding while Fc-enhancing variants at most, only moderately increased binding to FCRL5.

### IgG1-E293R selectively abrogates FCRL5-binding while preserving classical Fc**_γ_**R-binding

The lack of effect of FcγR-silencing mutations on FCRL5 binding suggests that the IgG residues involved in the interaction are distinct from the classical FcγRs. To further characterize the IgG-determinants that are essential in the interaction with FCRL5, we used a panel of IgG1-Fc mutants (From here on referred to as “T3-mutants”) that carry combinations of upper-CH2 to lower-CH3-spanning human-to-murine mutations of surface-exposed amino acids, previously used to map binding epitopes of rheumatoid factors to human IgG Fc (**Figure 4A**)^24^. The specific T3-mutant that carries a complete set of 20 CH2-CH3 mutations, T3-Bare, showed abrogated binding to FCRL5. Moreover, all T3-mutants that harbored upper-CH2 mutants showed abrogated binding compared to WT, whereas T3-mutants that harbored elbow-region or tail-region mutations were able to bind to FCRL5 D1-3. Using the T3-mutants BV7 and ER, we narrowed the essential FCRL5 binding residues to one or more of Y278, R292 and E293. FcγRIIb, which was taken along as a comparator, showed similar results (**Supplementary Figure 3A**). To assess which of these amino acids were involved in FCRL5 binding, we further assessed the binding to IgG1-mutants carrying either an R292W or E293R mutation (**Figure 4B**). The R292W mutation reduced binding to FcγRIIB, as reported previously^27^. Interestingly, FCRL5 binding was not affected by the R292W mutant, but was strongly reduced by the E293R mutant. SPR analysis confirmed the strong selectivity of the E293R mutant to abrogate binding to FCRL5, but not to FcγRIIA 131R, FcγRIIA 131H, FcγRIIB and FcγRIIIB (**Figure 4C**). Even FcγRIIIA 158F, which was reported to be sensitive to E293 mutations^27^, showed only a minimal reduction in affinity (1.5-2 fold). Taken together, our data show that E293 is a key residue in FCRL5 binding and that E293R selectively abrogates FCRL5 binding.

**Figure 4:**
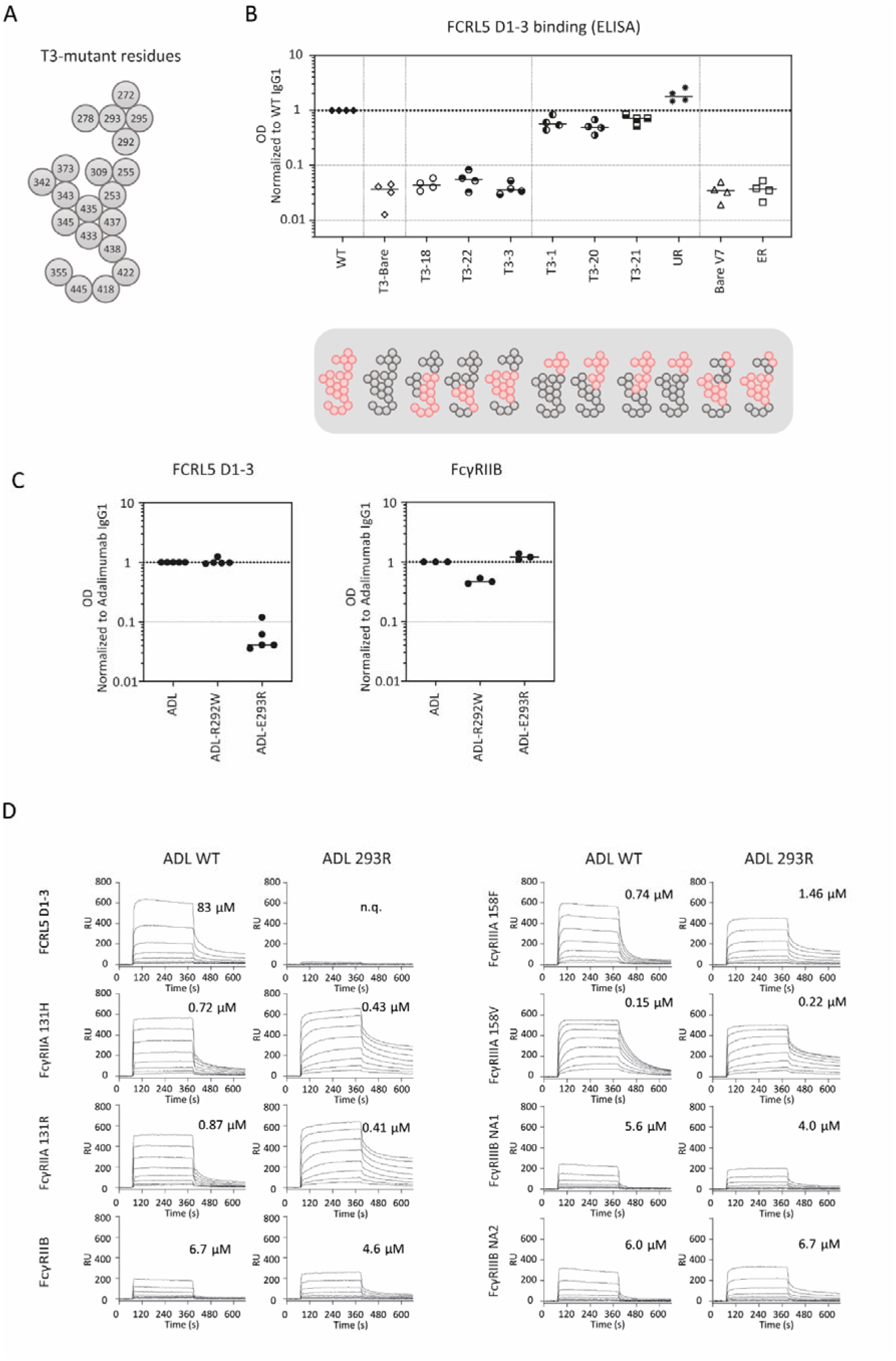
Binding of IgG-Fc mutants to FCRL5-D1-D3 and other classical Fc_γ_Rs receptors. (A) Visual representation of the location of the T3-mutants in the IgG Fc-tail. (B) ELISA binding of the T3-mutant to FCRL5 D1-3 (n=3). ODs are normalized to a-BT WT IgG1. (C) ELISA Binding of IgG1-R292W and IgG1-E293R to FCRL5 D1-3 (n=3). ODs are normalized to Adalimumab (IgG1). (D) Representative SPR sensorgrams of Adalimumab IgG1 and Adalimumab IgG1-E293R binding to FCRL5 and other classical FcγRs. The highest concentration of analyte here is 10 µM. KD calculations for low affinity receptors (FCRL5, FcγRIIB, FcγRIIIB) were performed with titrations starting at 100 µM.

### FCRL5 is a fucose-sensitive Fc receptor

IgG1-N297A was the only Fc-silenced IgG variant that largely abrogated binding to FCRL5, hinting to a potential role of the N297 IgG Fc glycan in the interaction with FCRL5. Loss of one of the IgG Fc glycan core-fucoses represents a physiologically relevant glycoform that has profound effects on binding to FcγRIIIA and FcγRIIIB^21,28^. Hence, we investigated if IgG1 fucosylation status could affect its binding to FCRL5.

We therefore measured FCRL5 binding to IgG-dimers from two clones on ELISA, comparing normal fucosylated IgG to low fucosylated IgG (**Figure 5A**). Interestingly, low fucosylated IgG1 showed a significant increase in FCRL5-binding potency compared to normal fucosylated IgG1, with a fold change of 4.5x. To further investigate this phenomenon, we compared the binding of FCRL5 D1-3 and D1-4 to low fucosylated IgG1-Fc and normal fucosylated IgG1-Fc by SPR (**Figure 5B**). Both FCRL5 D1-3 and D1-4 consistently showed a two- to three-fold increase in affinity to low fucosylated IgG1-Fc. Intrigued by these findings, we assessed the binding of low fucosylated IgG1 to FCRL5 using FCRL5-expressing HEK293T (FCRL5-HEK) cells. Binding was assessed for IgG as monomers (**Figure 5C**), or as dimers (**Figure 5D**). Up to five- to seven-fold enhanced binding was observed of the low fucosylated IgG1 upon incubating with FCRL5-expressing HEK293T cells. Physiologic IgG-immune complexes likely consist of more than two IgG molecules per complex. Moreover, small differences in affinity may result in large differences in binding in multivalent contexts, due to avidity effects. Therefore, we also assessed the effect of low fucosylated multivalent IgG-complexes on FCRL5 binding. To this end, we opsonized biotinylated human serum albumin (HSA-BT) with low or normal fucosylated anti-biotin IgG1 (**Figure 5E**). Again, we observed substantially increased binding of complexes with low fucosylated IgG to the FCRL5-expressing HEK293T cells. Taken together, FCRL5 preferentially binds afucosylated IgG over fucosylated IgG.

**Figure 5.**
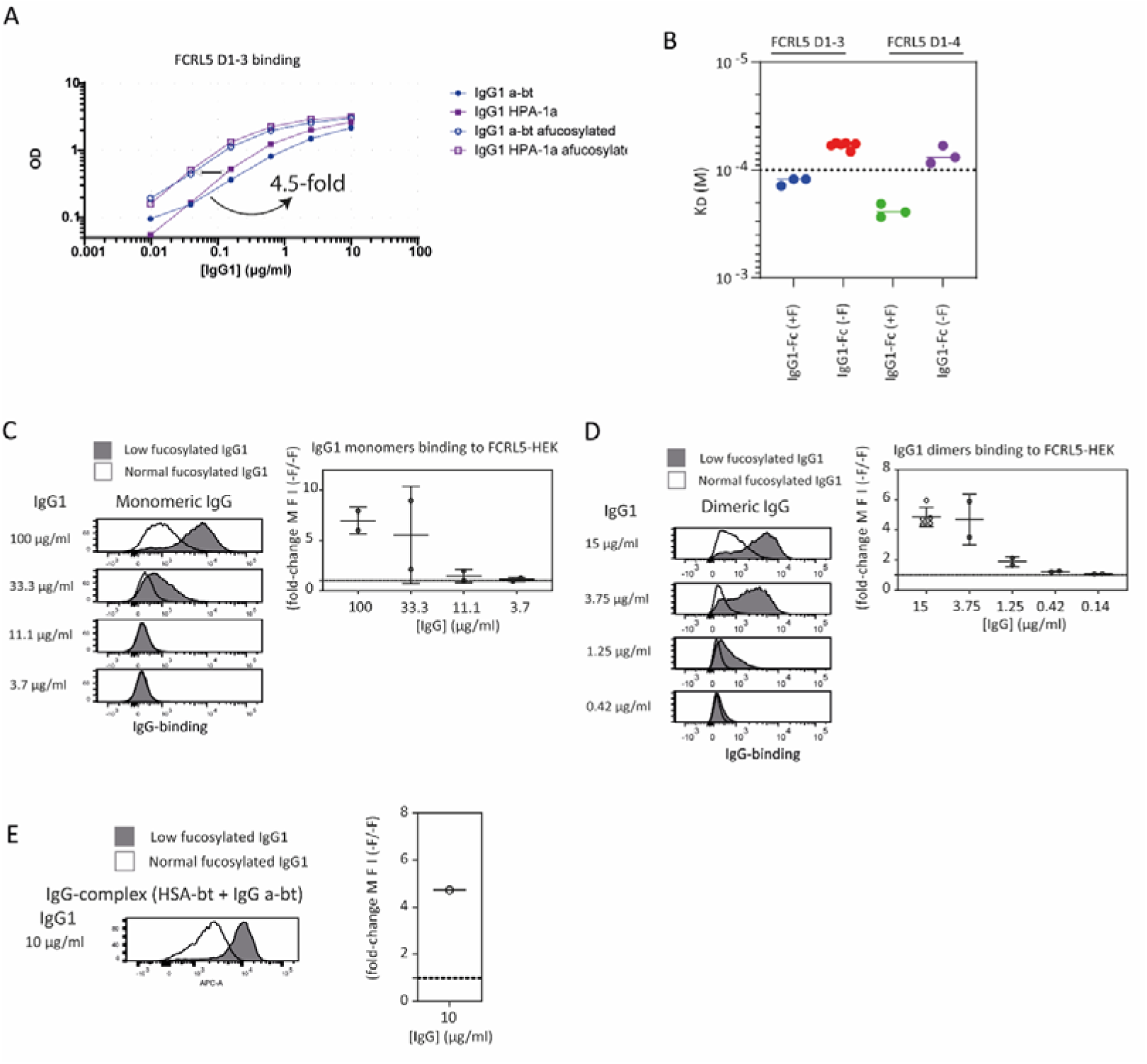
FCRL5 is a IgG Fc fucose-sensitive receptor. (A Binding of low and normal fucosylated IgG1 (anti-biotin and anti-HPA1a) to FCRL5 D1-3. (B) Graph depicting the calculated affinities of FCRL5 D1-3 and FCRL5 D1-4 for low and normal fucosylated IgG1-Fc. The dotted line represents the highest concentration of analyte measured (100 µM). (C) Flow cytometry histograms showing the binding of monomeric low or normal fucosylated IgG1 to FCRL5-HEK cells represented as median fluorescent intensity (MFI). Data from (N=2) experiments is normalized to normal fucosylated IgG1. (D) Flow cytometry histograms showing the binding of dimeric low or normal fucosylated IgG1 to FCRL5-HEK cells, represented as MFI. Data from (N=2) experiments is normalized to normal fucosylated IgG1. (E) Flow cytometry histograms showing the binding of HSA-BT opsonized with low or normal fucosylated IgG1 to FCRL5-HEK cells represented as MFI. Data is normalized to normal fucosylated IgG1.

## Discussion

Using complementary HP-SEC and SPR analyses, we demonstrate that FCRL5 is a bona fide IgG Fc-receptor that binds with very low affinity in a manner unique to FcγRs. We show that the first two N-terminal domains are sufficient to engage IgG, although the third N-terminal domain is required for maximal binding. Using an avidity-based ELISA, we show that many common Fc-silencing are not silencing for FCRL5 binding, and that Fc-enhancing IgG Fc-mutants do not significantly impact FCRL5 binding. Moreover, we show that E293R is a mutation that selectively abrogates FCRL5 binding, while preserving FcγR binding. Finally, we show that FCRL5 is a fucose-sensitive receptor that shows increased binding to afucosylated IgG1, as evidenced by ELISA, SPR and cellular flow cytometry assays.

FCRL5 has been previously described as a low-affinity IgG-receptor, however, reported affinities vary widely (∼1–45 µM)^16,17^. Such low affinities require careful consideration of experimental design and robust quality-checks of both the receptor and ligand, as high ligand concentrations are required for accurate affinity determinations to match the low K_D_. Moreover, ligand aggregates may more easily mask true K_D_ values for low-affinity interactions by disproportionate binding of aggregates due to avidity effects. Our data shows that FCRL5 has a K_D_ for IgG1 around 60-80 µM, slightly below 100 µM IgG, the highest concentration used in the SPR. These affinities place FCRL5 10 to 65 times lower in affinity for IgG1 than FcγRIIB and FcγRIIA, respectively.

Contrary to previously reported, our data do not support the notion that FCRL5 requires intact IgG for maximal binding. We observe similar affinities for FCRL5 binding to IgG1-Fc as we do for intact IgG1. Furthermore, we do not detect any binding of FCRL5 D1-3 to IgG1-F(ab’)_2_. These data are in favor of FCRL5 being an IgG Fc-receptor, like the classical FcγRs.

While truncated variants of FCRL5 containing the first three or four N-terminal domains (D1-3, D1-4), recapitulated binding of full-length, FCRL5 D1-9, the first two N-terminal domains (D1-2) were sufficient for IgG binding, as evidenced by the SPR sensorgrams. Its affinity for IgG1 however, was too low to be quantified accurately, suggesting that the true affinity for FCRL5 D1-D2 is likely around or below 100 µM. Of note, FcγRI, the sole FcγR with three domains, also requires the third domain for maximal IgG binding, which has been attributed to its role as a spacer that allows for the spatial accommodation of IgG, rather than direct interactions with IgG^29,30^. A cryo-EM structure of FCRL5 and hexameric IgG that was recently published, suggests that the third N-terminal domain of FCRL5 may interact with the CH3-domains of adjacent IgG molecules, thereby effectively bridging two IgG molecules in the hexamer structure^17^. These data would imply a reduction of binding to FCRL5 D1-2 via loss of interactions with this adjacent IgG CH3-domain. However, we do not have indications for higher-order interactions. An alternative explanation, also supported by this EM structure, is that intramolecular interactions between FCRL5 D1 and D3 stabilize FCRL5 in a quaternary structure optimal for IgG binding. A recent preprint by Herpers et al.^31^ that describes the cryo-EM structure of FCRL5 and a binding-enhanced IgG mutant, report a 1:1 interaction that does not involve engagement of an adjacent IgG Fc, further supporting this notion.

Our ELISA screen further indicates that two mutations in the upper CH3-domain, Q342L and Y373N do not affect binding to FCRL5, which is in contrast with the results of Chen et al. that these residues are directly involved in the interaction between the FCRL5 D2 and the IgG CH3-domain^32^. However, Herpers et al. report a modest decrease in affinity for replacement of Y373A. Similarly, the Q342L mutation in our T3-mutant panel may result in an affinity enhancement of similar magnitude, thereby countering the effects of Y373N.

Classical FcγRs display distinct binding preferences for different IgG subclasses. While the activating FcγR receptors show a clear preference for IgG1 and IgG3, the inhibiting receptor FcγRIIB shows comparable affinities for IgG1, IgG3 and IgG4. Similarly, our FCRL5 SPR measurements demonstrate strongest binding to IgG1, IgG3 and IgG4, and showed a three- to four-times lower affinity for IgG2, whereas the ELISA data indicate lower avidity for IgG3-containing immune complexes. However, due to minor fraction of aggregates in the IgG3 samples and the potentially distinct stoichiometry of IgG3-containing dimers resulting from its unique hinge architecture, the IgG3 data should be interpreted with caution. Therefore, we do not draw strong conclusions regarding subclass-specific FCRL5 interactions involving IgG3.

Modulating binding to FcγRs has been a key focus since the development of therapeutic mAbs, and multiple (glycoengineered) Fc-silencing and Fc-enhancing variants have been developed and implemented in IgG therapeutics^33^. Our data show FCRL5 is a bona fide Fc receptor, prompting us to assess binding of various Fc-modulating variants to FCRL5. We find that most Fc-variants designed to prevent FcγR interactions, do not prevent FCRL5 binding. This means that FCRL5 may be (co-)engaged in settings where it may not be appropriate or wanted. For example, obexelimab, a CD19/FcγRIIB bi-specific Ab that is used for B cell inhibition in autoimmune settings^34^, may co-engage FCRL5 on B cells that co-express FcγRIIB and FCRL5. Due to the lack of knowledge on the biological function of FCRL5, however, it is unclear what unintended effects may arise from such interactions. We find that the E293R mutation can be exploited to differentiate in binding mode between FCRL5 and the classical FcγRs. Namely, this mutation abrogates binding to FCRL5 while maintaining binding to FcγRIIA, FcγRIIB, FcγRIIIA and FcγRIIIB. Furthermore, E293R may be combined with existing Fc-modulating formats to further fine-tune specific receptor targeting. Utilization of E293R and LALAPG may further help elucidate the relative contributions of FCRL5 and FcγRIIB to IgG-mediated B cell modulation in experimental settings, and IgG1-SELF, an FcγRII-enhanced variant that is known to increase affinity for FcγRIIB up to 430-fold^35^ but does not affect affinity for FCRL5, may be of additional interest.

Glycoengineering represents another important strategy for modulating Ab affinity for FcγRs^21–23^. In particular, due to the increased affinity of afucosylated IgG for FcγRIII, IgG afucosylation is a physiologically relevant phenotype which is observed in various protective immune responses and pathogenic (allo)immune responses^21,36–40^. We demonstrate that FCRL5 also shows a modest increase in affinity for afucosylated IgG, of which multivalent contexts reveal large physiological differences in binding, as evidenced by flow cytometry on FCRL5-expressing HEK cells. As FCRL5 is considered a hallmark of DN2 B cells ^5,6^, afucosylated IgG immune complexes may allow for DN2 B cell-specific feedback. Hence, further research into the association of FCRL5-expressing B cells and afucosylated IgG responses is needed.

Concluding, we provide a comprehensive analysis on the FCRL5-IgG interaction that define FCRL5 as a very low affinity, bona fide Fc receptor with a preference for afucosylated IgG.

## Materials and methods

### Protein production and purification

#### Commercial recombinant FCRL5 and Fc_γ_Rs

Commercial recombinant human FCRL5 (FC5-H82E3) (AA: 16-851) was bought from Acro Biosystems. Biotinylated human CD32B/FCGR2B His & AviTag (10259-H27H-B), Biotinylated human CD32A/FCGR2A 167H His & AviTag (10374-H27H1-B), Biotinylated human CD32A/FCGR2A 167R His & AviTag (10374-H27H-B), Biotinylated CD16A/FCGR3A 176V His & AviTag (10259-H27H1-B), Biotinylated CD16A/FCGR3A 176F His & AviTag (10259-H27H-B) were bought from Sino Biologics.

#### Therapeutic monoclonal antibodies

Therapeutic monoclonal antibodies were obtained from commercial autoinjectors. Prior to affinity measurements, the mAbs were rebuffered from their original storage buffer to PBS using centrifugal filters (Amicon MWCO, Merck).

#### Fc-engineered IgG monoclonal antibodies

A panel of Fc-engineered anti-CD20 antibody heavy-chain expression vectors were constructed by inserting de novo synthesized (Geneart) codon optimized HC coding regions into expression vector pcDNA3.3 (Invitrogen). The HC coding regions consisted of the VH regions of human mAbs 7D8 (human CD20-specific [Teeling et al 2004]) genetically fused to the CH regions of human IgG1*03 or one of the mutant variants: (EU numbering conventions are used throughout the manuscript):

- FcγR-silenced: LALA (L234A, L235A)^41^, PA (P329A)^42^, PG (P329G)^41^, LALAPA (L234A, L235A, P329A)^41^, LALAPG (L234A, L235A, P329G)^41^, ΔG236 (deletion of G236)^43^, N297A^44^, IgG1-lower IgG2 hinge (IgG1-lh2; E233P, L234V, L235A, ΔG236)^45^
- FcγR-enhanced: DLE (S239D, A330L, I332E)^46^, DFTE (S239D, S267E, H268F, S324T)^47^, EFT (S267E, H268F, S324T)^47^, SELF (S276E, L328F)^48^
- FcRn-silenced: AARA (I253A, H310A, H435A)^49^ (and K409R to enable controlled Fab-arm exchange)^50^
- FcRn-enhanced: DQ (T256D, T307Q)^51^

Likewise, separate light-chain expression vector were constructed by inserting the 7D8 VL coding regions in frame with the CL coding regions of the human (J00241) kappa light chain into expression vector pcDNA3.3.

All antibodies were produced under serum-free conditions by co-transfecting relevant heavy and light chain expression vectors in FreeStyle™ Expi293F™ cells, using ExpiFectamine™ 293 (LifeTechnologies), according to the manufacturer’s instructions.

IgG1 antibodies were purified by protein A affinity chromatography (MabSelect SuRe; GE Health Care), dialyzed overnight to PBS and filter-sterilized over 0.2-µM dead-end filters. Alternatively, IgG3 antibodies were purified by protein G affinity chromatography (GE Health Care). Purity was determined by CE-SDS and concentration was measured by absorbance at 280 nm (specific extinction coefficients were calculated for each protein). Batches of purified antibody were tested by high-performance size-exclusion chromatography (HP-SEC) for aggregates or degradation products and shown to be at least 95% monomeric. Purified antibodies were stored at 2-8°C.

#### In-house production of recombinant proteins

In-house FCRL5 (D1-2, D1-3, D1-4 and D1-9), IgG mAbs (including (therapeutic) mAbs expressed as Fc-mutants or other subclasses), murine anti-human kappa light chain IgG (clone K35) and rat anti-mouse kappa light chain Fab-CH1 (clone RM19) were produced by transfection of HEK293F cells with pcDNA3.1 or pEE6.4 plasmids encoding the respective proteins. FCRL5 (D1-2, D1-3, D1-4 and D1-9) were expressed with a C-terminal 10-Histag and AviTag. RM19 Fab-CH1 was expressed with a C-terminal AviTag.

Transfections were performed as previously reported^52^. In short, 100 million HEK293F cells were transfected with 31 µg pSVLT/p21/p27 plasmid was co-transfected with 34 µg FCRL5 plasmid using PEI-Max (Polysciences, 24765) and Opti-MEM (ThermoFisher Scientific, 31985070). 4-7 days post-transfection, cells were spun down (10-30 minutes, 1000-2000g) and supernatant was removed and filtered through .45 µm filters (Whatman, FP30/0.45 CA-S). Proteins were subsequently purified with the ÄKTA Start chromatography system (see below). Site-specific biotinylation of proteins was performed using Enzymatic Protein Biotinylation Kit (CS0008, Sigma-Aldrich).

#### Proteolytic-generation of murine anti-human kappa light chain F(ab’)_2_ (K35)

Murine K35 IgG was rebuffered 3-times, from PBS to 0.1M sodium acetate (pH 3.7) using centrifugal filters (Merck, 10 kDa MWCO). Prior to rebuffering, the columns were equilibrated with 0.1M sodium acetate (pH 3.7).

Pepsin (Sigma-Aldrich, Cat. No. P6887) was equilibrated to room temperature before opening and added to the IgG sample at a 1:100 (w/w) ratio (pepsin:IgG). Samples were incubated at 37 °C for 2.5 h.

The enzymatic reaction was irreversibly terminated by bringing up the pH to 8.5-9 with 1M Tris-HCl (pH 9.0). The sample was subsequently simultaneously depleted from digested Fc peptides and enriched for F(ab‘)2 fragments using a sequential anti-murine light chain column and a multispecies Fc-directed column (manufacturer and column information mentioned below under Protein affinity purification). F(ab‘)2 fragments were eluted with 0.1M glycine (pH 3.0) and immediately neutralized to pH 7.5 with 2M Tris-HCl. F(ab‘)2 fragments were finally rebuffered to PBS using centrifugal filters (Merck, 10 kDa MWCO) and verified by SDS-PAGE.

#### Protein affinity purification

In-house produced proteins were purified using the ÄKTA Start chromatography system (Cytiva) with Cytiva affinity purification columns (HisTrap HP, HiTrap Prot G HP, HiTrap Prot A HP) or in-house poured columns CaptureSelect IgG-Fc multispecies affinity matrix (2942852050, ThermoFisher Scientific) and CaptureSelect KappaXP affinity matrix (2943212005, ThermoFisher Scientific) and CaptureSelect LC-kappa murine affinity matrix (191315010, ThermoFisher Scientific). Peak fractions were collected and rebuffered to PBS or HEPES + 10% trehalose (FCRL5) or 10 mM acetate buffer pH 4.5 (antibodies), sterile filtered, and stored at -80°C and -20°C, respectively.

### FCRL5 and Fc**_γ_**RIIB binding experiments

#### FCRL5 and Fc_γ_RIIB IgG binding ELISA

Biotinylated FCRL5 and FcγRIIB were incubated for 2 hours at a concentration at 1 µg/mL in 100 µl PBS per well in 96-well streptavidin-coated plates (Pierce, Ref: 15500). IgG was incubated with murine anti-human kappa light chain F(ab‘)2 K35 in a 1:1.1 (w/w) ratio of IgG:K35 for 1 hour in PTG buffer at RT while shaking. Next, the streptavidin-coated plate was washed 5 times with wash buffer (0.2% Tween-20 in PBS). After incubation, the IgG:K35 complexes were diluted and subsequently transferred to the streptavidin-coated plate. Samples were incubated for 1 hour at RT while shaking and subsequently washed 5 times with wash buffer. IgG:K35 complexes were detected by incubation with a rat anti-murine kappa light chain F(ab) HRP at 0.5 µg/mL in PTG for 30 minutes at RT, while shaking. TMB (1-Step Ultra-TMB, ThermoFisher, No. 34029) was used to visualize the binding of IgG:K35 complexes to FCRL5 and FcγRIIB. The TMB reaction was stopped with with 100 µL of 0.2 M sulfuric acid (HLSOL). Lastly, ODs were measured at 540nm-450nm on a Millipore BioTek 800 TS microplate reader.

#### FCRL5 and Fc_γ_R affinity measurements

FCRL5 and FcγR affinity measurements were performed using the IBIS MX96 (IBIS Technologies) system and an IBIS Array Printer (IBIS Technologies). C-terminally biotinylated FCRL5 or hFcγRs at 5 3-fold titrated concentrations were captured simultaneously on a SensEye G-streptavidin sensor (Ssens; cat#1-09-04-006) in PBS (Fresenius Kabi; cat#X0022-145) containing 0.075% (v/v) Tween-80 (Merck; cat#P4780). The starting concentration for the FCRL5 variants ranged from 60 nM to 10 nM and was 10 nM for FcγRIIA 131R and FcγRIIB. Next, a 2-fold dilution series of the analyte was injected ranging from 0.781 μM to 100 μM in PBS (Fresenius Kabi) at pH 7.4 supplemented with 0.075% (v/v) Tween-80 (Merck). Regeneration of the sensor surface between the cycles was performed by injection of 10 mM Glycine-HCl pH 3.0 (Merck; cat#1.00590 – Merck; cat#1.00317). Affinity (KD) calculations were performed using an equilibrium analysis, interpolating to an Rmax of 500 response units (RU). A 1:1 Langmuir binding model was used for fitting, assuming equilibrium to be reached after 360s of IgG containing analyte injections, as described previously^53^. Analysis and calculations were performed using Scrubber Software Version 2 (BioLogic Software), Excel (Microsoft) and Graphpad Prism (Prism).

#### FCRL5 IgG binding cellular experiments

The binding of IgG to FCRL5 was assessed on HEK293T cells expressing FCRL5. Monomeric IgG was titrated from a starting concentration of 100 µg/mL. To generate dimeric IgG complexes, human IgG was incubated with mouse anti-human kappa F(ab’)L (clone K35) at a w/w ratio of 1:0.9 (IgG:K35) with a starting IgG concentration of 15 µg/mL To generate opsonized HSA-BT particles, human IgG was incubated with HSA-BT and anti-biotin IgG at a w/w ratio of 1:5 (IgG:HSA-BT), at an IgG concentration of 10 µg/mL. All complexes were pre-incubated for 1 hour prior to use.

700.000 HEK293T WT or 500.000 HEK293T-FCRL5 cells were seeded per well in U-bottom 96-well plates, centrifuged at 200 × g for 4 min, and washed twice with FACS buffer (PBS, 1% w/v bovine serum albumin). Cells were incubated with the pre-incubated IgG complexes for 1 hour at 4 °C in the dark, followed by two times washing with FACS buffer. Detection was performed by adding 100 µL APC-conjugated anti-human kappa Fab (MH19 Fab) for monomeric and HSA-BT-based complexes, or APC-conjugated anti-mouse kappa Fab (RM19 Fab) for dimeric complexes, and incubating for 30 minutes at 4 °C in the dark. Cells were washed another two times with FACS buffer. Samples were analyzed using a BD LSR II flow cytometer.

### Cell culture and generation on FCR5L-expressing HEK293T cells

#### Cell culturing

Human embryonic kidney 293T (HEK293T) cells were cultured in Dulbecco’s Modified Eagle Medium (DMEM) with GlutaMAX™ (Gibco, Cat. No. 61965-026) supplemented with 1% penicillin–streptomycin (100 U/mL penicillin and 100 µg/mL streptomycin) at 37 °C in a humidified incubator with 5% CO□.

Stable FCRL5-expressing HEK293T cells were cultured in DMEM with GlutaMAX (Gibco, Cat. No. 61965-026) supplemented with 10% fetal bovine serum, 1% penicillin–streptomycin, and 5 µg/mL blasticidin (InvivoGen, Cat. No. ant-bl-1) at 37 °C in a humidified incubator with 5% CO□.

### Protein quality assessment

#### HPLC-SEC

Proteins were fractionated using HP-SEC Agilent 1260 Infinity II (Agilent Technologies) with a Superdex 200 Increase SEC column 10/300 GL (Cytiva) and PBS as running buffer. Elution was monitored by measuring the absorption at 280 nm. The molecular weight of peaks was determined from monitoring multiangle laser light scattering (MALS) using a mini-DAWN (Wyatt Technology Europe) and refractive index using an Optilab (Wyatt Technology Europe).

#### Data analysis

Flow cytometry data were analyzed using FlowJo v10.10.0.

