## supplementary data for "FCRL5 is a fucose-sensitive IgG-Fc receptor with binding properties distinct from classical Fcγ receptors"

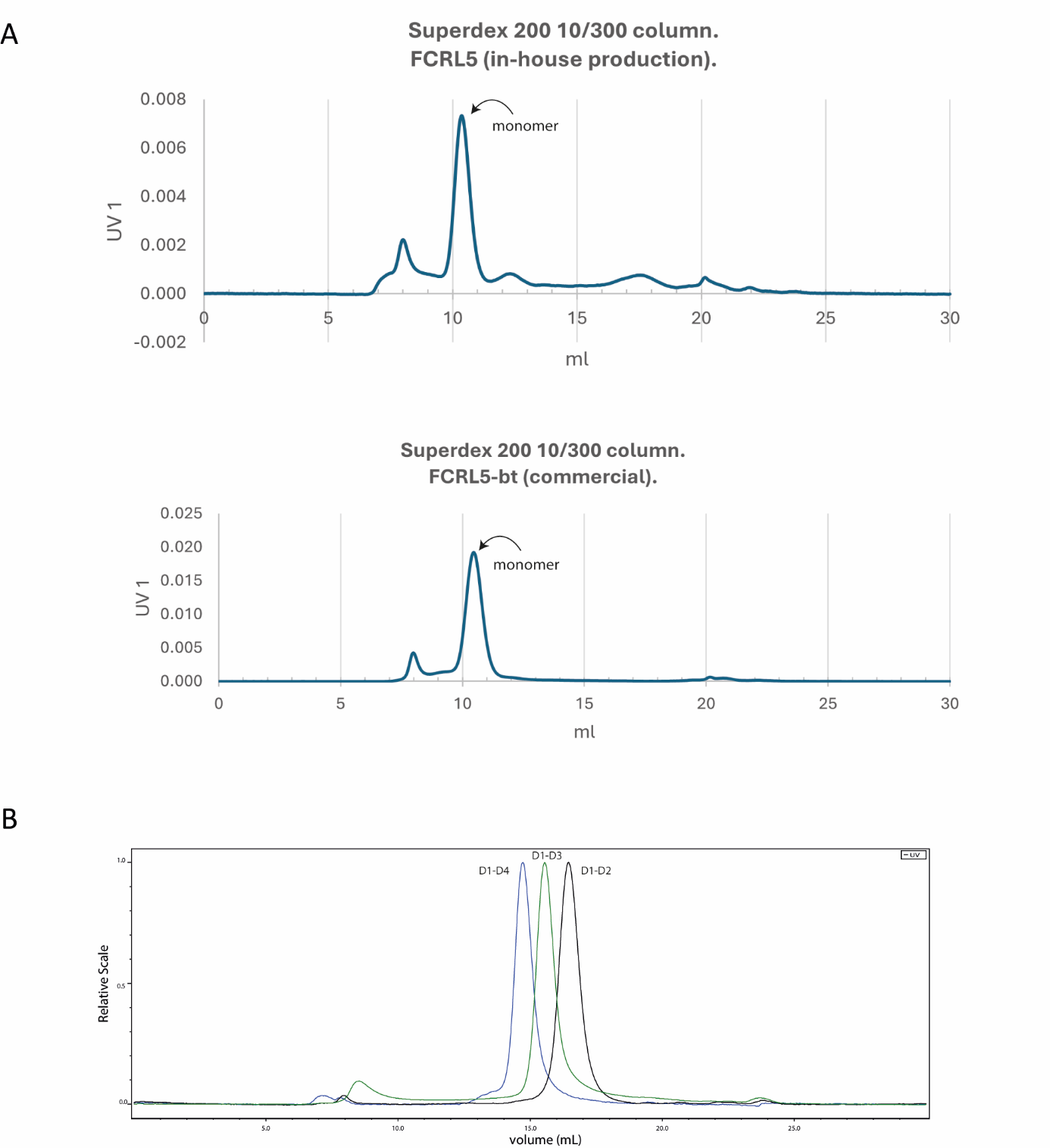

**Supplementary Figure 1: HPLC-SEC analysis of commercial and in-house produced FCRL5 variants.**

(A) Graph depicting the UV absorption at 280nm elution profile of full length D1-D9 FCRL5 (in-house produced and commercial) using the HP-SEC Agilent 1260 Infinity II with a Superdex 200 Increase SEC column 10/300 GL (Cytiva)

(B) Graph depicting the UV absorption at 280nm elution profile of truncated FCRL5 variants (D1-D2, D1-D3 and D1-D4) using the HP-SEC Agilent 1260 Infinity II with a Superdex 200 Increase SEC column 10/300 GL (Cytiva)

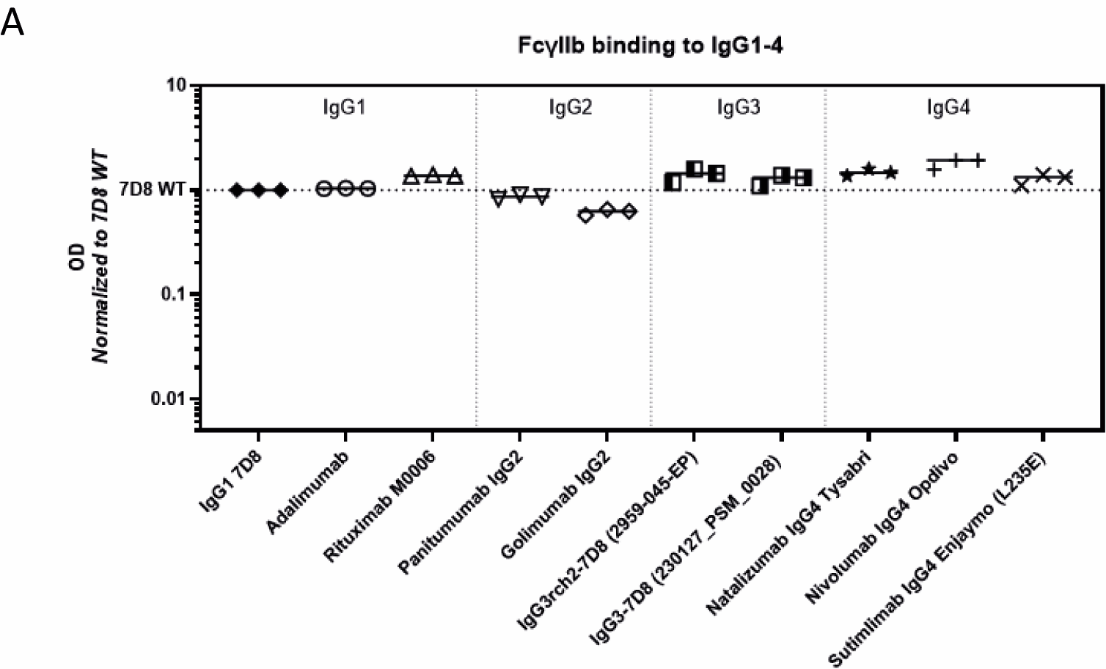

**Supplementary Figure 2: Avidity-driven ELISA setup measuring binding of IgG subclasses to FcγRIIB.**

(A) Binding of various IgG1-4 mAbs to FcγRIIB (n=3). ODs are normalized to 7D8-IgG1.

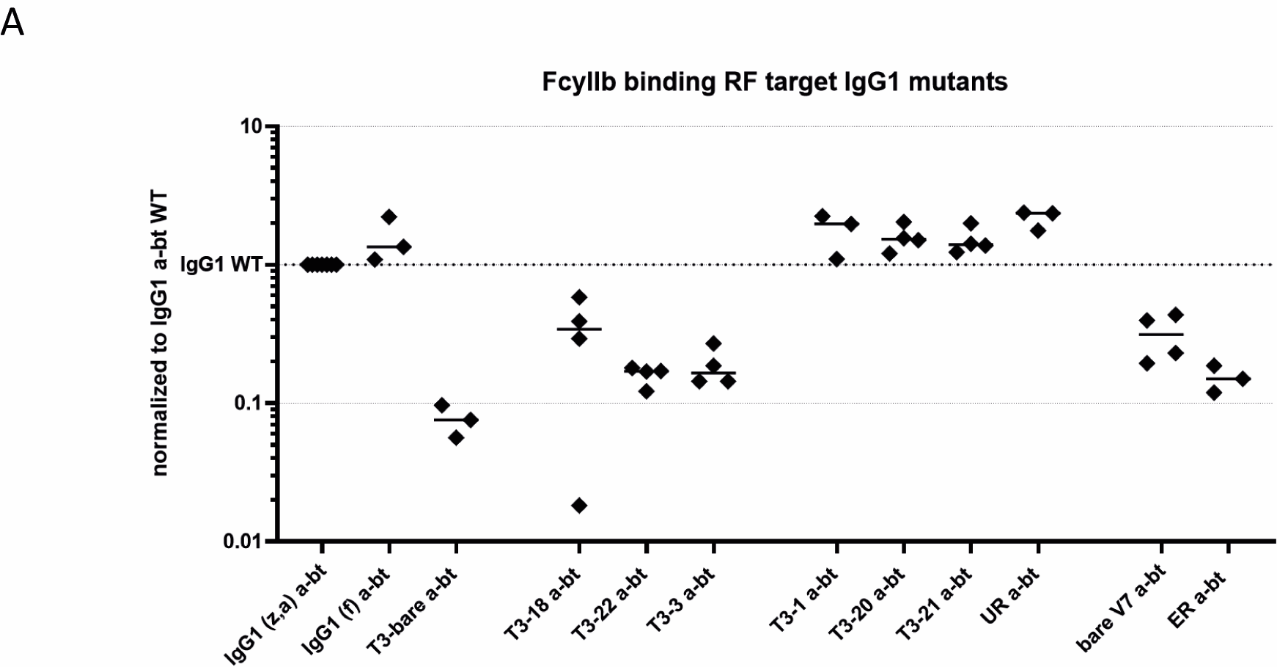

**Supplementary Figure 3: Binding of IgG-Fc mutants to FcγRIIB.**

(A) ELISA binding of the T3-mutant to FcγRIIB (n=3). ODs are normalized to a-BT WT IgG1.
